# A Small interfering RNA lead targeting RNA-dependent RNA-polymerase effectively inhibit the SARS-CoV-2 infection in Golden Syrian hamster and Rhesus macaque

**DOI:** 10.1101/2020.07.07.190967

**Authors:** Se Hun Gu, Chi Ho Yu, Youngjo Song, Na Young Kim, Euni Sim, Jun Young Choi, Dong Hyun Song, Gyeung Haeng Hur, Young Kee Shin, Seong Tae Jeong

## Abstract

A small interfering RNA (siRNA) inhibitors have demonstrated the novel modality for suppressing infectious diseases. Sixty-one siRNA molecules, predicted by the bioinformatics programs, were screened for the possibility of treating severe acute respiratory syndrome coronavirus 2 (SARS-CoV-2) using an *in vitro* plaque assay. Among six siRNA leads with the efficacy of reducing plaque number, the siRNA targeting RNA-dependent RNA polymerase (RdRp) showed a reduction in SARS-CoV-2 infection-induced fever and virus titer in the Golden Syrian hamster and rhesus macaque. These results suggest the potential for RdRp targeting siRNA as a new treatment for the coronavirus disease 2019 (COVID-19).

## Introduction

Coronaviruses (family *Coronaviridae*) are enveloped viruses with a positive-sense and single-stranded RNA genome. Some coronaviruses cause diseases of varying clinical severity, such as severe acute respiratory syndrome (SARS) and the Middle East respiratory syndrome (MERS) (1). More recently, SARS coronavirus 2 (SARS-CoV-2), which caused the coronavirus disease 2019 (COVID-19) pandemic, has infected approximately 11,301,850 people and caused 531,806 deaths (as of July 6, 2020) globally (2). Despite the severity of the COVID-19 pandemic, there are no specific drugs except for a comprehensive treatment and management guide, which consist of symptomatic treatment, supportive therapy, and/or antiviral/antibiotic therapy. However, the comprehensive therapy shows different clinical outcomes for each patient because the efficacy and effectiveness of the therapy depends on the medical status of each patient. Current trials of repurposing of drugs for COVID-19 have shown no acceptable risk-benefit ratios except remdesivir and dexamethasone. However, there are several candidate drugs for COVID-19 treatment under research or clinical trials, such as antiretroviral drugs (HIV-1 protease inhibitor), RNA-dependent RNA polymerase (RdRp) inhibitors (remdesivir, favipiravir), antiviral cytokines (interferon β), and anti-spike protein monoclonal antibodies. Despite emergency use authorization of remdesivir for the treatment of severe COVID-19 patients, no drugs have been approved for COVID-19 so far.

RNA interference (RNAi) has a specific mechanism to silence gene expression by degrading messenger RNA (mRNA) targeted by small RNA molecules, such as microRNAs (miRNAs) and small interfering RNAs (siRNAs), which are complementary to mRNA (3, 4). Recently, it has been shown that RNAi specifically silences viral gene expression and treats infectious diseases caused by a viral infection (5, 6). Because RNAi treatments regulate gene expression inducing diseases, they have the advantage of being used to develop drugs against infectious diseases that are difficult to treat using chemical or small-molecule compounds (6). Moreover, RNAi is an adequate method to deal with a pandemic because RNAi can be synthesized and tested in a very short time when the target sequence is identified.

Our goals in this study are to develop siRNAs to target and silence viral genes of SARS-CoV-2 for the inhibition of viral replication and treatment of COVID-19. In addition, since siRNAs can be used for versatile treatment against other types of coronavirus infections, the siRNAs were further selected by matching the target sequences in the highly conserved regions with SARS-CoV-1 or SARS-CoV-2 variants. It is expected to reduce the risk of ineffectiveness due to mutation at the target sequence (7). Among putative 61 siRNA sequences targeting the conserved sequences of structure and replication genes, we obtained six siRNA leads that showed a reduction in cytopathic effect (CPE) and the plaque assay *in vitro*. Finally, we chose one of the most effective siRNAs that was less affected by mutation, and then performed *in vivo* experiments with siRNA in the Syrian hamster and rhesus macaque to confirm its protective efficacy against SARS-CoV-2.

This study indicates that siRNAs are a rapid and effective treatment for new emerging pathogens such as SARS-CoV-2, which may be able to diminish the impact of possible future pandemics by developing treatments promptly.

## Materials and Methods

### Virus and cell culture

SARS-CoV-2 isolated from a COVID-19 patient in Korea was used. The pathogen resources (NCCP43326) for this study were provided by the National Culture Collection for Pathogens (8). Viruses were propagated in Vero E6 cells (CRL 1586, American Type Culture Collection, Manassas, VA, USA) and maintained in Dulbecco’s modified Eagle’s medium (DMEM) supplemented with 2% heat-inactivated fetal bovine serum (FBS; GIBCO, USA), 2 mM L-glutamine, and antibiotics (penicillin/streptomycin) at 37°C for 3 days in a 5% CO_2_ incubator. As determined using plaque assay, the infectivity titers of the SARS-CoV-2 stocks were 2 × 10^6^ plaque-forming units (PFU/mL).

### Design of siRNA targeting SARS-CoV-2

In this study, we obtained the SARS-CoV-2 genome sequence of MT039890 from (https://www.ncbi.nlm.nih.gov/nuccore/MT039890). We focused on seven regions of the SARS-CoV-2 genome, namely the leader sequence, replicase polyprotein1a (pp1a), RdRp, spike protein (S), nucleocapsid protein (N), membrane protein (M), and envelope protein (E) to design specific siRNAs to inhibit viral replication. Potential siRNA sequences targeting viral genes were predicted using the programs siDirect (http://sidirect2.rnai.jp/) and VIRsiRNApred (http://crdd.osdd.net/servers/virsirnapred/). Among the siRNAs predicted, we examined whether siRNA predicted sequences that targeted the sequence of SARS-CoV, to identify the 21 siRNA sequences targeting the conserved region in coronaviruses. We further selected 40 siRNA sequences from the rest of the predicted siRNAs that did not match SARS-CoV according to two criteria: first, siRNAs that have a score of less than off-target effects in two portals; second, siRNAs that have low mutation rate among variants from Nextstrain (https://nextstrain.org/). We finally obtained 61 putative siRNAs, all of which were 21-mer with dTdT at the 3’-overhang. The siRNAs were synthesized by Bioneer Co. (Daejeon, Republic of Korea).

### *In vitro* efficacy test using Vero E6 cells (siRNA transfection)

Vero E6 cells were inoculated into a 24-well plate, 1 mL each, through a 10% FBS DMEM without penicillin/streptomycin to ensure that the 24-well plate is 80–90% confluent. The next day, the Vero E6 cells were transfected with 100 nM siRNA using Lipofectamine RNAiMAX transfection reagent (Invitrogen, USA), according to the manufacturer’s instructions. The cells were incubated at 37°C in 5% CO_2_ for 3 h and washed with phosphate-buffered saline. Next, the SARS-CoV-2 stock of 1 × 10^5^ PFU/mL was diluted at 1/100 into 1×10^3^ PFU/mL. Next, 200 µL each was added to make the final 200 PFU. The Vero E6 cells were then infected with SARS-CoV-2 for 1 h 30 min. After infection, the supernatant was removed, changed with 2% FBS DMEM medium, and incubated at 37°C for 3 days in a 5% CO_2_ incubator. The cells were observed daily for CPE using a microscope (Zeiss, Germany).

### *In vitro* efficacy test using Vero E6 cells (plaque assay)

To assess viral titers, a plaque assay was performed using Vero E6 cells in 6-well culture plates. Briefly, subconfluent monolayers of Vero E6 cells were inoculated with 10-fold serial diluents and incubated at 37°C for 1 h 30 min in a 5% CO_2_ incubator. After incubation, the supernatant was removed and carefully overlaid with 1 mL/well of overlay solution (1:2 mixtures of 1% agarose and 2% FBS DMEM) and incubated for 3 days. The plates were then fixed and inactivated using a 4% formaldehyde solution and stained with 0.1% crystal violet.

#### RNA extraction and cDNA synthesis

Total RNA was extracted from supernatants and animal tissues using the PureLink RNA mini kit (Invitrogen, San Diego, CA, USA) according to the manufacturer’s instructions. cDNA was synthesized using the SuperScript III First-Strand Synthesis Systems (Invitrogen) and then analyzed for SARS-CoV-2 RNA using qRT-PCR.

### *In vitro* efficacy test using Vero E6 cells

The primers and probe targeting the SARS-CoV-2 E gene and the forward and reverse primer sequences for real-time PCR were 5’-ACAGGTACGTTAATAGTTAATAGCGT-3’ and 5’-ATATTGCAGCAGTACGCACACA-3’, respectively, and the probe was 5’-ACACTAGCCATCCTTACTGCGCTTCG-3’ (9). The probe was labeled with the reporter dye 6-carboxyfluorescein at the 5’-end and quencher dye Black Hole Quencher 1 at the 3’-end, respectively. Each 20-μL reaction mixture contained 2 μL cDNA, 10 μL 2X TaqMan Gene Expression master mix (Applied Biosystems, USA), 0.5 μL forward and reverse primers (36 μM), 0.5 μL fluorescent probe (10 μM), and 6.5 μL double deionized water. The reaction was performed at 50°C for 2 min and 90°C for 10 min, followed by 40 cycles at 95°C for 15 s and 60°C for 1 min, in a QuantStudio 6 Flex Real-Time PCR system (Applied Biosystems, USA). The standard curve was constructed using RNA from SARS-CoV-2 infected Vero E6 cells. RNA concentration was measured using a NanoDrop spectrophotometer. After RT-PCR using random primers (10 μM), 10-fold serial dilutions of the cDNA were used in duplicate to generate a standard curve.

### Animal experiments

We assessed the therapeutic efficacy of siRNA in the Syrian hamster and rhesus macaque and established a model for SARS-CoV-2 infection (10, 11). First, we confirmed whether Syrian hamsters were infected with SARS-CoV-2; 6-week-old male Syrian hamsters were randomly segregated into 5 groups. Each group of the animals was inoculated with 4-40,000 PFU of SARS-CoV-2 via the intranasal route. At 2 days post-infection (d.p.i.), the lungs were harvested for qRT-PCR analysis. Next, we evaluated the therapeutic efficacy of siRNA in the Syrian hamsters. SARS-CoV-2 inoculation of the animals was performed through intranasal instillation of 1,000 PFU per head. Four hours after virus inoculation, the animals in groups 2 and 3 were intranasally administered with siRNA (No. 14) 17.3 μg (low dose) and 34.6 μg (high dose), respectively (Table 2). At 2 d.p.i., the lungs were harvested for further analysis.

**Table 1.**
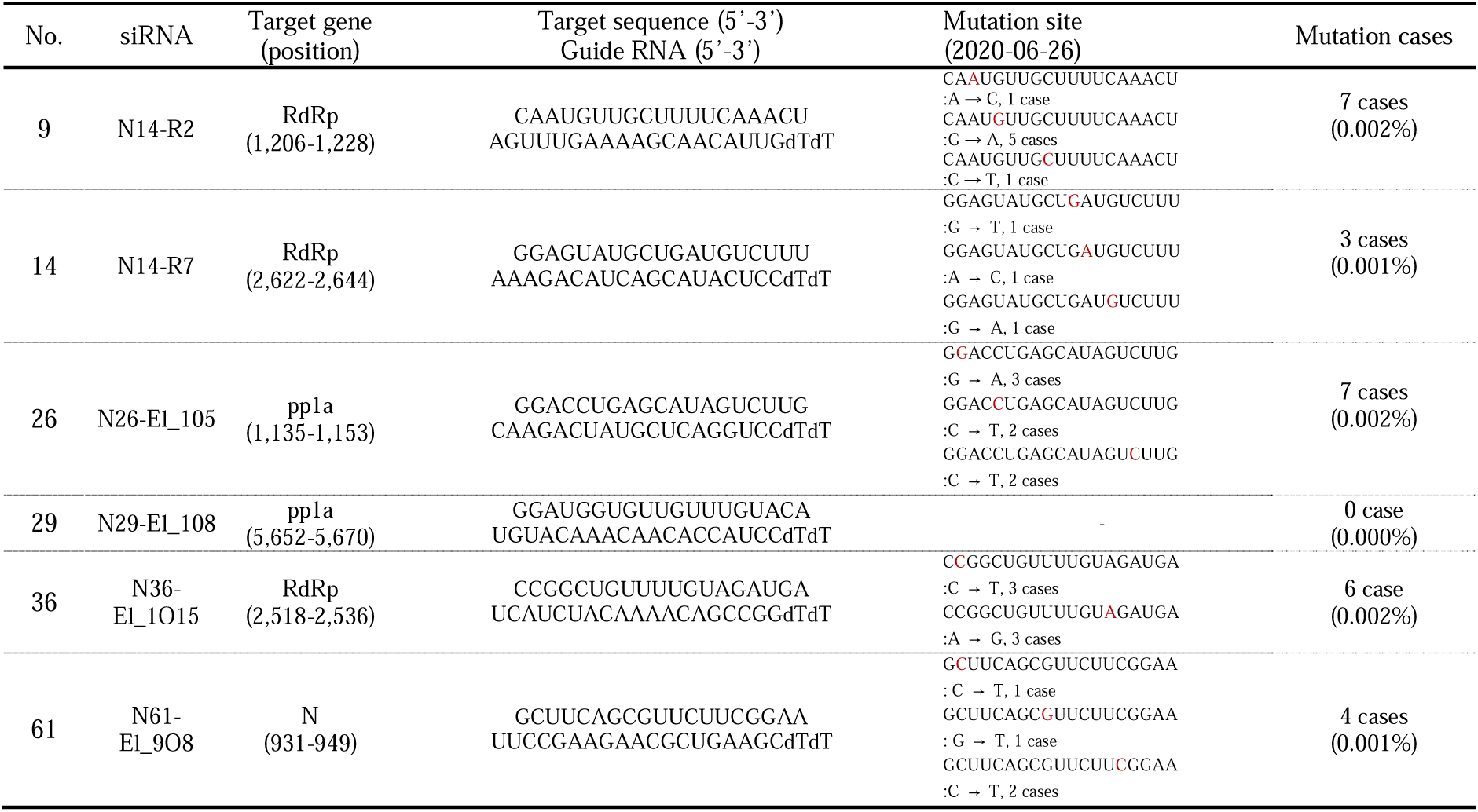
The sequences of siRNA targeting SARS-CoV-2.

**Table 2.** Study design of Syrian Hamsters.

| Group | Sex | Age<br>(weeks) | No. of<br>Animals | Virus<br>inoculation | siRNA<br>Dose | Dose<br>regimen |
| --- | --- | --- | --- | --- | --- | --- |
| G1 | Male | 6 | 5 | §IN, 1.0x10<sup>3</sup><br>PFU/head | 0 | - |
| G2 | Male | 6 | 5 | §IN, 1.0x10<sup>3</sup><br>PFU/head | 17.3ug/head | +4h |
| G3 | Male | 6 | 5 | §IN, 1.0x10<sup>3</sup><br>PFU/head | 34.6ug/head | +4h |

§ IN, Intra Nasal

Third, we assessed the therapeutic efficacy in the rhesus macaque. Three male rhesus macaques were randomly segregated into three groups of one each (Table 3). The animals were inoculated with 4.0 × 10^6^ PFU of SARS-CoV-2 through intranasal and intratracheal routes under anesthesia. After 4 and 24 h, the animals in groups 2 and 3 were intratracheally administered with siRNA (No.14) 2 mg/kg (low dose) and 4 mg/kg (high dose), respectively, under anesthesia. All the animals were monitored daily for clinical signs, body weight, and body temperature. Swab samples, including nasal, oropharyngeal, and rectal swabs, were collected. The animals were euthanized at 3 d.p.i. All inoculations and handling of the animals, as well as the method of euthanasia and collection of tissues, were performed according to well-established protocols approved by the Institutional Animal Care and Use Committee of the Agency for Defense Development (ADD-IACUC-20-12 and ADD-IACUC-20-13). All experiments were conducted in consultation with the veterinary and animal care staff of the ADD animal biosafety level-3 (ABSL-3) containment, in a facility in which other respiratory disease-causing coronaviruses had never been handled.

**Table 3.** Study design of rhesus macaques.

| Group | Sex | Age<br>(years) | Body<br>weight<br>(kg) | Virus<br>inoculation | siRNA<br>Dose | Dose<br>regimen | Total<br>dosage |
| --- | --- | --- | --- | --- | --- | --- | --- |
| G1 | Male | 5 | 5.4 | ¶IN, IT,<br>4.0x10<sup>6</sup><br>PFU/head | 0 | - | - |
| G2 | Male | 4 | 3.5 | ¶IN, IT,<br>4.0x10<sup>6</sup><br>PFU/head | 2mg/kg | +4h, +24h | 14mg |
| G3 | Male | 4 | 3.0 | ¶IN, IT,<br>4.0x10<sup>6</sup><br>PFU/head | 4mg/kg | +4h, +24h | 24mg |

¶ IN, Intra Nasal. IT, Intra Trachea

## Results

### Design of siRNA targeting SARS-CoV-2

To design a siRNA targeting SARS-CoV-2, siRNA molecules were screened from the whole sequence of SARS-CoV-2 (SNU-MT039890). We designed siRNA sequences and then selected a siRNA that targeted the conserved sequence of SARS-CoV-2 for application in the treatment of various strains of SARS-CoV-2. To target the S proteins, we designed siRNAs in the less mutated region (HR2) and receptor binding motif (RBM). Among sixty one siRNA sequences expected to inhibit SARS-CoV-2 infection, twenty one siRNAs, targeting leader sequence, RdRp, S, and N, were primarily selected as the conserved region sequences of coronavirus. Additionally, forty siRNA sequences, targeting pp1a, RdRp, S, N, E, and M, were designed to be most conserved for SARS-CoV-2 and its mutants. All siRNA candidates were 21-mer by adding two deoxythymidine (dTdT) at the 3’ overhang to enhance siRNA stability and binding efficiency to target RNA (12).

### Selection of efficient siRNA leads using *in vitro* efficacy test

We tested the efficacy of siRNAs to inhibit viral infection in the Vero E6 cells, in which the CPE can be confirmed by apoptosis upon SARS-CoV-2 infection and plaque reduction assay (13). After the Vero E6 cells were infected with 200 PFU of SARS-CoV-2 and treated with 100 nM of the 61 siRNA sequences, the CPE and plaque assays were performed at 3 d.p.i. (Fig. 1a-1c). At 3 d.p.i., all siRNA candidates reduced CPE in SARS-CoV-2 infected Vero E6 cells, while CPE was observed in the control group (data not shown). Based on the results of the CPE and plaque assays, we selected six siRNA leads, showing a reduction in plaque number. Six siRNA leads were subjected to the NCBI blast to ensure that they did not target any human genes. RdRp, pp1a, and N mRNAs were targeted by 3 siRNAs (Nos. 9, 14, and 36), 2 siRNAs (No. 26 and 29), and 1 siRNA (No. 61), respectively.

**Fig. 1.**
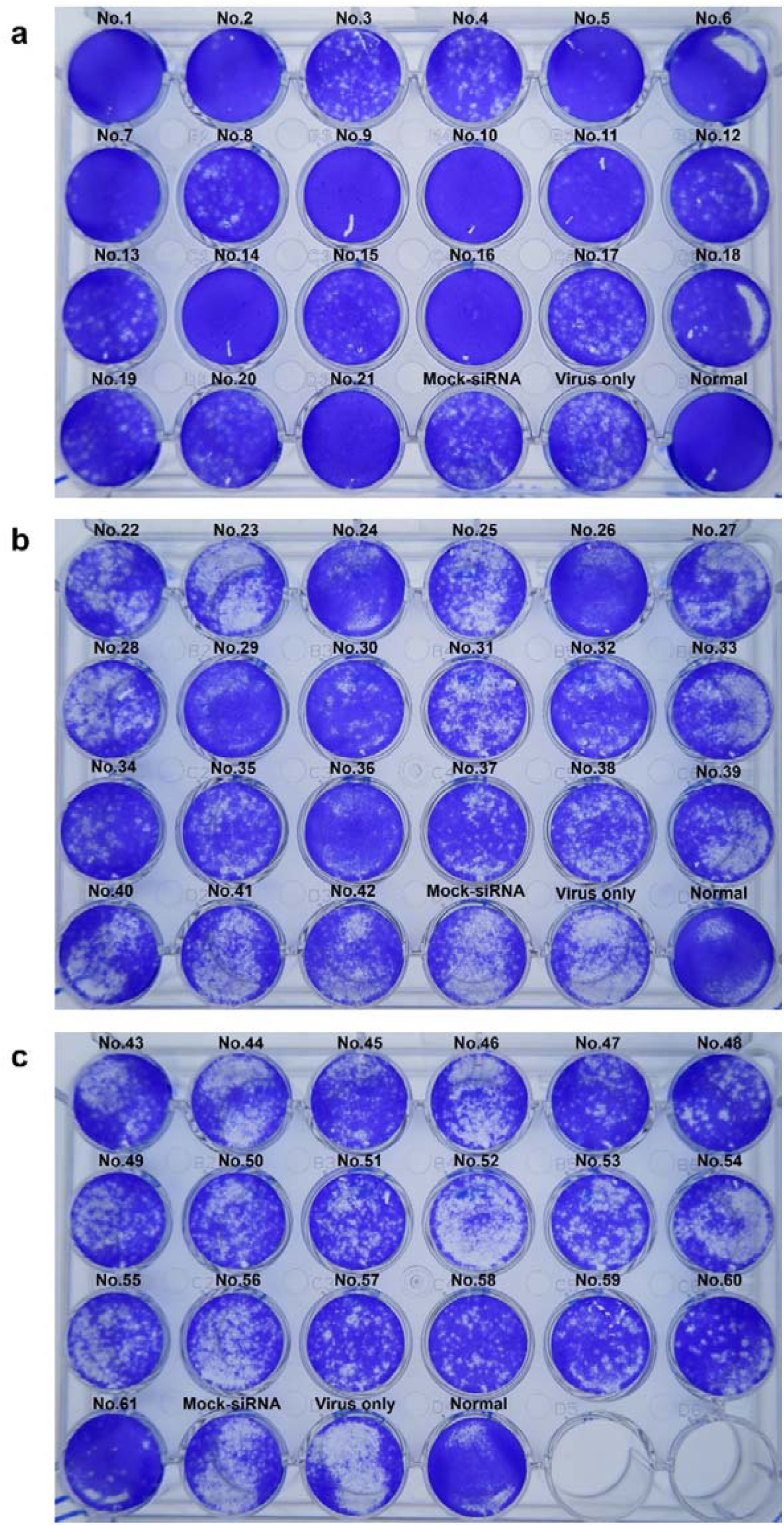
Screening of siRNA molecules for SARS-CoV-2 using plaque assay. (a) siRNA No. 1-21. (b) siRNA No. 22-42. (c) siRNA No. 43-61.

The Nextstrain website was used to monitor the data on SARS-CoV-2 mutation and to confirm whether the regions targeted by our siRNA leads were less mutated (as of June 26, 2020). Mutation rates in siRNA-targeted regions were 0-0.002%, indicating that siRNA leads can be used to treat most SARS-CoV-2 strains. Hence, the six siRNA leads were effective in reducing viral replication *in vitro* and less influenced by SARS-CoV-2 mutation (Table 1).

### Efficacy test of the No. 14 siRNA lead against SARS-CoV-2

With six siRNAs, we examined whether the replication of SARS-CoV-2 was inhibited using a plaque reduction assay. Vero E6 cells were infected with 500 PFU of SARS-CoV-2 with or without siRNA candidates and the plaque assay was performed at 3 d.p.i. From the results, lead siRNA No. 14 (out of the six candidates) showed the least plaque formation (data not shown). To calculate the half-maximal effective concentration (EC_50_), the Vero E6 cells infected with 500 PFU of SARS-CoV-2 were treated with the siRNA No. 14 in a dose-dependent manner (5-100 nM) (Fig. 2a-2i). Then, the CPE assay was performed at different time points. The CPE started to reduce with 20 nM of the siRNA No. 14, and cell morphology was similar in cells treated with 100 nM of the siRNA No. 14. EC_50_ was further calculated using qRT-PCR to determine the copy number of SARS-CoV-2 (Fig. 2m). From the result, EC_50_ of the siRNA No. 14 was approximately 9.7 nM at 2 d.p.i. in 500 PFU SARS-CoV-2 infected Vero E6 cells. Recently, it was reported that EC_50_ of the antiviral drugs, remdesivir, chloroquine and nafamostat, known to be effective for COVID-19 (14).

**Fig. 2.**
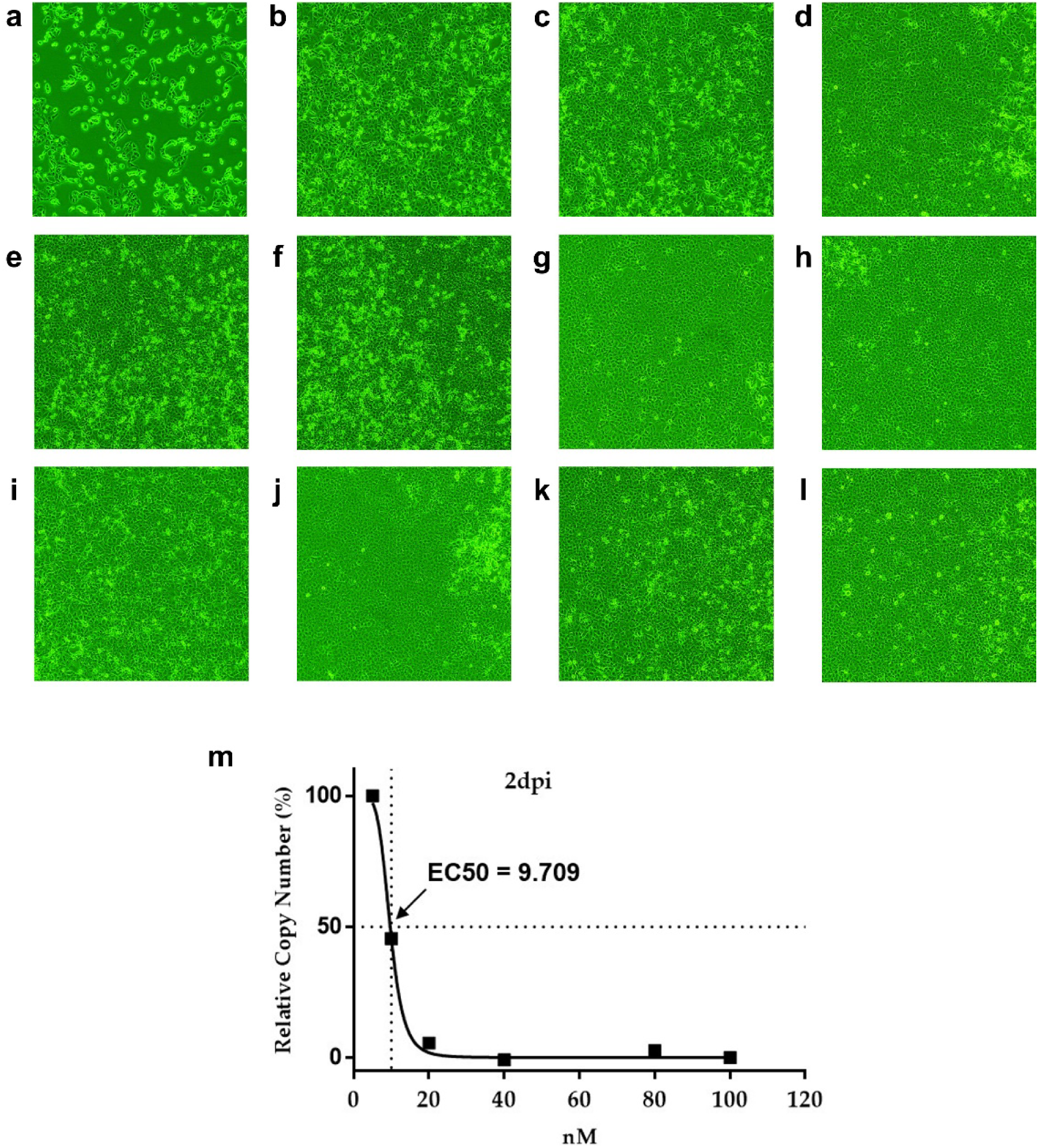
siRNA No. 14 inhibition of SARS-CoV-2 cytopathicity in Vero E6 cells (CPE assay and EC_50_). Vero E6 cells were infected with SARS-CoV-2 and incubated for 2 days. (a) Mock-siRNA (100 nM). (b) 5 nM. (c) 10 nM. (d) 20 nM. (e) 30 nM. (f) 40 nM. (g) 50 nM. (h) 60 nM. (i) 70 nM. (j) 80 nM. (k) 90 nM. (l) 100 nM. (m) EC_50_ of siRNA No. 14 using qRT-PCR.

### Clinical signs in golden Syrian hamsters and rhesus macaques

To assess the therapeutic efficacy of siRNA in the hamsters, three groups were formed composed of a control group and two treatment groups (Table 2). At 2 d.p.i., the control hamsters showed a hunched posture, ruffled hair, and mild cough. However, clinical signs such as cough were not observed in the siRNA No. 14-treated hamsters. The three rhesus macaques consisted of one control animal and two treatment animals (Table 3). After anesthetizing the animals, three male rhesus macaques were inoculated with 4 × 10^6^ PFU SARS-CoV-2, administered at a dose of 5 mL through intratracheal (4 mL) and intranasal (1 mL) instillation. After 4 and 24 h, we administered the siRNA No. 14 through the same route at low (2 mg/kg; G2) and high (4 mg/kg; G3) doses, respectively. At 1 d.p.i., diarrhea was observed in the viral control and body temperature was markedly elevated compared to the base level (Fig. 3c). SARS-CoV-2-infected rhesus macaques (G1) had elevated body temperature, from 38.6-40.4°C, between 1 and 2 d.p.i. Interestingly, high-dose siRNA No. 14-treated animals (G3) showed a base level of body temperature over 3 days. All treated animals displayed normal appetite and behavior.

**Fig. 3.**
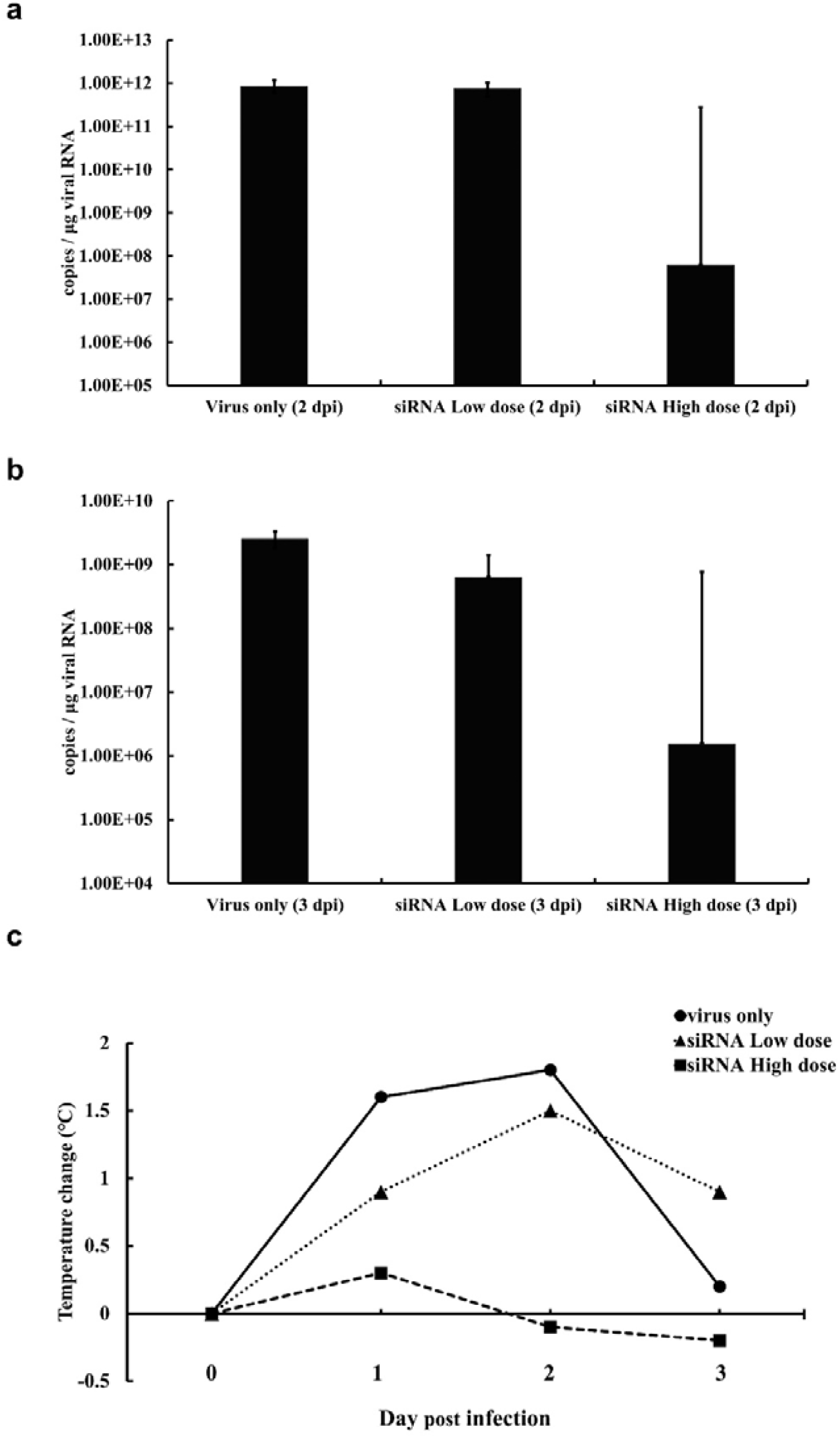
qRT-PCR of viral RNA copies in lung of Syrian hamsters and trachea of rhesus macaques inoculated with SARS-CoV-2, respectively. Viral RNA copies per 1 μg of total RNA are sacrificed on 2 and 3 d.p.i. (a) Syrian hamsters. (b) Rhesus macaques. (c) Body temperature change of rhesus macaques.

### qRT-PCR analysis of the Syrian hamsters and rhesus macaques

To confirm infection of SARS-CoV-2 in the hamster, we inoculated each group with 4-40,000 PFU SARS-CoV-2 by the intranasal route. SARS-CoV-2 RNA was detected in all hamsters, including the 4 PFU inoculation group (data not shown). We assessed the therapeutic efficacy of the siRNA No. 14 in the hamsters and analyzed the lung of the hamsters at 2 d.p.i. using qRT-PCR. The results showed that the number of viral RNA copies from the lung tissue of the siRNA No. 14-treated hamsters markedly decreased by approximately 10^4^ viral copies compared to that of the control hamsters (Fig. 3a). Next, we investigated viral copies from the trachea of rhesus macaques at 3 d.p.i. using qRT-PCR. The number of viral RNA copies in the siRNA No. 14-treated rhesus macaques significantly decreased by approximately 10^3^ viral copies compared to that in the control animals (Fig. 3b). These results indicate that siRNA No. 14 effectively inhibited SARS-CoV-2 infection and replication in the lung and upper respiratory tract.

## Discussion

An effective SARS-CoV-2 treatment is needed to end the COVID-19 pandemic in the near future. To date, many reports have demonstrated that RNAi is a powerful method for gene silencing, including virus infection and replication *in vitro, in silico* and *in vivo* (15, 16, 19) against SARS-CoV. Wang et al presented that SARS-CoV replication was efficiently inhibited by siRNA in Vero cells. Li et al demonstrated that siRNA was significantly effective in prophylactic and therapeutic regimen against SARS-CoV in Rhesus macaques (19). More recently, a CRISPR/Cas13d system was reported for the therapeutic agent of SARS-CoV-2 (20). In addition, Chen et al showed theoretical predictions of the potential siRNA targets by computation modeling in the SARS-CoV-2.

Animal studies on SARS-CoV-2 play an important role in understanding the pathogenesis and development of therapeutic drugs. Recently, animal studies on SARS-CoV-2 have reported that rhesus monkeys and transgenic mice expressing human ACE2 receptor were susceptible to SARS-CoV-2 infection (17, 18). In addition, the pathogenesis and transmission of SARS-CoV-2 were shown in Syrian hamsters (10). Efficacy test using siRNA therapeutics for SARS-CoV-2 has been reported not yet in the hamster and rhesus macaque. However, efficacy tests including DNA vaccines and therapeutic antibody were reported in the SARS-CoV-2 animal models (21, 22). The Syrian hamster model was used for demonstrating protective efficacy of neutralizing antibodies against SARS-CoV-2. The hamsters showed typical clinical signs within one week after virus inoculation. Rhesus macaque model was used for DNA vaccine test. Rhesus monkeys developed humoral and cellular immune responses and, were protected against SARS-CoV-2.

In this study, to demonstrate the siRNAs that are effective for the inhibition of SARS-CoV-2 infection, we tested 61 siRNA duplexes in Vero E6 cells. Among them, the best lead, siRNA No. 14, showed strong inhibitory efficacy against SARS-CoV-2 infection and replication in Vero E6 cells. To assess the inhibition efficacy of siRNA No. 14 for SARS-CoV-2 in animals, we used Syrian hamsters and rhesus macaques. Our data showed that SARS-CoV-2 viral RNA decreased in the siRNA No. 14-treated animals. Additional research is warranted to determine the possibility of siRNA treatment for COVID-19. Further studies would also be needed to address the safety and effective delivery methods of siRNA.

## Acknowledgements

This work was supported by grants (011555-912664201) from the Agency for Defense Development, Republic of Korea.

## Author Contributions

S.H.G., C.H.Y. Y.K.S. and S.T.J. conceived and designed the experiments; S.H.G., C.H.Y. and Y.S. performed the experiments; S.H.G., C.H.Y., Y.S., N.Y.K., E.S., J.Y.C., D.H.S., G.H.H., Y.K.S. and S.T.J. analyzed the data and prepared the figures; and all authors prepared and reviewed the manuscript.

## Competing interests

The authors have no conflicts of interest.

## Notes

### Competing Interest Statement

The authors have declared no competing interest.

